# Identification of cytoplasmic chaperone networks relevant for respiratory syncytial virus replication

**DOI:** 10.1101/2022.02.23.481726

**Authors:** Victor Latorre, Ron Geller

**Author notes:** **Corresponding author:** Ron Geller.

## Abstract

RNA viruses have limited coding capacity and must therefore successfully subvert cellular processes to facilitate their replication. A fundamental challenge faced by both viruses and their hosts is the ability to achieve the correct folding and assembly of their proteome while avoiding misfolding and aggregation. In cells, this process is facilitated by numerous chaperone systems together with a large number of co-chaperones. In this work, we set out to define the chaperones and co-chaperones involved in the replication of respiratory syncytial virus (RSV). Using an RNAi screen, we identify multiple members of cellular protein folding networks whose knockdown alters RSV replication. The reduced number of chaperones and co-chaperones identified in this work can facilitate the unmasking of specific chaperone subnetworks required for distinct steps of the RSV life cycle and identifies new potential targets for antiviral therapy. Indeed, we show that the pharmacological inhibition of one of the genes identified in the RNAi screen, valosin-containing protein (VCP/p97), can impede the replication of RSV by interfering with the infection cycle at multiple steps.

## INTRODUCTION

The majority of RNA viruses have small genomes and encode only few proteins. Nevertheless, RNA viruses are able to perform an astonishing number of functions with their limited proteome, including disarming antiviral defenses, replicating the viral genome, producing new virions, and spreading to new cells and/or hosts. To a large part, this is achieved by masterfully interacting with, and coopting, cellular processes to favor viral replication at each step of the virus life cycle.

Among the key challenges faced by viruses and cells alike is the successful translation of their genome to a functional proteome. In cells, this process is facilitated by a large number of dedicated protein-folding factors, termed chaperones, that help maintain cellular homeostasis (proteostasis) together with the protein degradation machinery (Kim *et al*., 2013; Balchin, Hayer-Hartl and Hartl, 2016). To perform their functions, chaperones are organized into networks that cooperate to support folding from the time the polypeptide emerges from the ribosome, through maturation and assembly into larger protein complexes, as well at later points, when proteins become misfolded or aggregated (Kim *et al*., 2013; Balchin, Hayer-Hartl and Hartl, 2016; Bar-Lavan, Shemesh and Ben-Zvi, 2016). The major chaperone systems in human cells are comprised of numerous isoforms of Hsp70/HSPA, a constitutive and a stress-isoforms of Hsp90/HSPC, and the oligomeric chaperone complex TRiC/CCT (Kim *et al*., 2013; Balchin, Hayer-Hartl and Hartl, 2016; Bar-Lavan, Shemesh and Ben-Zvi, 2016; Radons, 2016b, 2016a). To facilitate client protein folding and maturation, these chaperones employ ATP-hydrolysis to drive conformational changes in their client proteins. These major chaperone systems are regulated by a much larger cohort of co-chaperones that provide client-protein specificity and interconnect different chaperone systems. For example, the binding of client proteins to Hsp70 and the chaperone cycle are regulated by >50 DNAJ/Hsp40 proteins and nucleotide exchange factors, such as BAG domain proteins (Kampinga and Craig, 2010; Bracher and Verghese, 2015). Similarly, Hsp90 activity is regulated by numerous co-chaperone that regulate client-protein binding and release (Schopf, Biebl and Buchner, 2017). Finally, the activity of both the Hsp70 and Hsp90 chaperones systems are linked via interaction with a large set of proteins containing tetratricopeptide repeat (TPR) domains (Balchin, Hayer-Hartl and Hartl, 2016). Aside from these central systems, a set of small heat shock proteins (HSPBs) form part of the cellular response to combat protein aggregation, and the prefoldin chaperone participates in early folding events en route to the central chaperone systems (Balchin, Hayer-Hartl and Hartl, 2016).

While chaperones are mainly involved in protein folding, they participate in the degradation of terminally misfolded proteins via both the proteasomal and autophagy pathways (Edkins, 2015; Ciechanover and Kwon, 2017). This is accomplished by co-chaperones that help link individual chaperones with the degradation machinery. For example, the C-terminus of Hsc70-interacting protein (CHIP) is an E3 ubiquitin ligase that ubiquitinates misfolded proteins to tag them for proteasomal degradation. Via its TPR domain, CHIP interacts with both Hsp70 and Hsp90, linking these chaperone systems to the cellular degradation machinery (Edkins, 2015; Dikic, 2017). Additionally, valosin-containing protein (VCP)/p97 is an ATP-dependent molecular multifunctional protein that assists in the remodeling of protein complexes as well as in the extraction of proteins across membranes, as in ER-associated degradation (ERAD) (Dikic, 2017; Ye *et al*., 2017; Huryn, Kornfilt and Wipf, 2020). While VCP/p97 is frequently involved in facilitating the degradation of proteins, it can also act to disassemble complexes (Ye *et al*., 2017).

As is the case with cellular proteins, the proteins of RNA viruses are folded by cellular chaperone systems. Indeed, pharmacological inhibition of Hsp70 and Hsp90 has revealed that many RNA viruses are not only dependent on these central chaperone systems for their replication but are in fact hypersensitive to changes in their levels compared to host cells (Mayer, 2005; Geller, Taguwa and Frydman, 2012; Wang *et al*., 2017; Aviner and Frydman, 2019). Accordingly, reductions in the activity of these chaperones at levels that are not toxic to cells are highly detrimental to viral replication, making chaperone inhibitors broad-spectrum antivirals (Mayer, 2005; Geller, Taguwa and Frydman, 2012; Wang *et al*., 2017; Aviner and Frydman, 2019). However, a more general understanding of chaperone networks utilized by particular viruses is largely lacking. Recently, different components of the Hsp70 chaperone system were shown to play distinct roles at different stages of the replication cycle of both Dengue virus and ZIKA virus (Taguwa *et al*., 2015, 2019).

In this work, we aimed to obtain a general picture of chaperones utilized by a medically relevant human RNA virus, respiratory syncytial virus (RSV). RSV is among the most important respiratory infections in the pediatric setting and causes significant morbidity in the elderly population (Griffiths, Drews and Marchant, 2017; Bergeron and Tripp, 2021). RSV belongs to the *Mononegavirales* order, harboring a single-stranded, negative-sense RNA genome, and encodes for 11 different proteins. Hence, RSV represents a relatively simple model with which to begin to untangle cellular protein folding networks. While prior work has shown Hsp70 and Hsp90 to play a key role in the replication of RSV (Radhakrishnan *et al*., 2010; Geller, Andino and Frydman, 2013; Munday *et al*., 2015a, 2015b; Huong, Tan and Sugrue, 2016; Latorre, Mattenberger and Geller, 2018), general information about additional chaperone systems and/or co-chaperones is lacking. To this end, we conducted an RNAi screen for a large number of cytoplasmic chaperones and co-chaperones in human cells. We identify numerous players in the protein-folding network to modulate RSV replication. This information can be used as a starting point to map specific chaperone subnetworks required for distinct steps of the RSV infection cycle. In addition, some of the identified factors can present potential targets for antiviral therapy. Indeed, we show that pharmacological inhibition of one of the identified genes, the VCP/p97 protein, can block the replication of RSV as well as an additional member of the *Mononegavirales*, supporting its potential as a broad-spectrum antiviral target.

## MATERIAL AND METHODS

### Cell lines, viruses, and reagents

HeLa-H1 (CRL-1958), HEp2 (CCL-23), HEK-293 (CRL-1573), and A549 (CCL-185) cells were obtained from ATCC. All cells were maintained in DMEM with 10% FBS, 100 IU/mL penicillin, and 100 μg/mL streptomycin at 5% CO_2_ and 37°C. RSV expressing the fluorescent protein mKate2 (RSV-mKate2) was generated by transfection of the infectious clone system obtained from BEI Resources (NIAID, NIH: Bacterial artificial chromosome plasmid pSynkRSV-I19F containing antigenomic cDNA from respiratory syncytial virus (RSV) A2-Line19F, NR-3646). This was done by transfection of HEK-293 cells with an optimized T7 plasmid (Addgene 65974) together with the antigenomic plasmid, N, P, M2-1, and L plasmids at a ratio of 4:2:2:2:1 using Lipofectamine 2000 (ThermoFisher Scientific). Cells were transfected in 6 well plates and subsequently transferred to T25 flasks with HEp2 cells until cytopathic effect (CPE) was observed. For the generation of RSV encoding nanoluciferase (RSV-nanoLuc), the mKate2 open reading frame was replaced with that of the nanoluciferase gene from pNL1.2[NlucP] (Promega) and the virus was generated as indicated above. Passage 0 stocks were amplified 3 additional times and titered using TCid50 (RSV-nanoLuc) and plaque assay (RSV-mKate2; see below). Vesicular stomatitis virus (VSV) encoding mCherrry was kindly provided by Dr. Rafael Sanjuan (University of Valencia). CVB3 was generated as previously described (Mattenberger *et al*., 2021). The VCP/p97 inhibitor CB-5083 (CAS: 1542705-92-9; Cayman Chemicals) and the hsp90 inhibitor 17-DMAG (CA 467214-20-6; LC Laboratories) were dissolved in DMSO and resazurin (CAS: 62758-13-8; Sigma-Aldrich) was dissolved in water.

### Virus quantification

Plaque assays were used to measure RSV, VSV, and CVB3 infectious virus production. For this, serial dilutions of the virus were used to infect confluent HeLa-H1 cells in 6 well plates for 2 hours (RSV-mKate2) or 45 minutes (VSV-mCherry and CVB3). Subsequently, the virus was removed and cells overlaid with DMEM containing 2% FBS and either 0.6% agarose for RSV-mKate2 and VSV-mCherry or 0.8% agarose for CVB3 during 24 hours. Finally, RSV-mKate2 plaques were assessed by examining fluorescence using a live-cell microscope (IncuCyte S3; Sartorius), while VSV-mCherry and CVB3 plaques were visualized using crystal violet staining following fixation with 1% formaldehyde. For the TCid50 assay, confluent Hep2 cells in a 96 well plate were infected with serial dilutions of RSV-nanoLuc (8 replicates) in DMEM containing 2% FBS. Seven days after the infection, cells were fixed with 1% formaldehyde and stained with crystal violet. To quantify intracellular virus replication, the mean fluorescent signal in each well resulting from the expression of the viral encoded fluorescent protein (mKate2 or mCherry) was obtained using a live-cell microscope.

### RNAi screening

Genes were selected for evaluation based on a list of the human chaperone network (Brehme *et al*., 2014), which was filtered to exclude non-cytoplasmic chaperones and those not expressed in A549 cells or lung tissue based on data from the human protein atlas (https://www.proteinatlas.org/). This list was inspected individually, and missing known chaperone/co-chaperone genes were included (Table S1). We then ordered endoribonuclease prepared siRNA (MISSION esiRNA, Sigma-Aldrich) for all genes for which esiRNA were available (Table S3). For the primary screen, A549 cells were reverse transfected with 20uM of esiRNA using the MISSION siRNA Transfection Reagent (Sigma-Aldrich) following the manufacturer protocol in white-bottom 96 well plates. After 48 hours, cells were infected with RSV-nanoLuc at a multiplicity of infection (MOI) of 0.01. Finally, RSV replication was evaluated by reading nano-luciferase activity with the Nano-Glo Luciferase Assay system (Promega) on a Tecan Spark plate reader. All experiments were performed in 6 replicates on 6 independent plates. The strictly standardized mean difference (SSMD) relative to infected wells that were transfected with a control esiRNA targeting firefly luciferase was calculated using the UMVUE estimate of SSMD (formula A5 in table 3 in (Zhang, 2011)). Finally, p-values were converted to q-values to correct for multiple testing using the R package q-value (version 3.14). For the secondary validation, esiRNAs targeting a different region of the gene were employed (esiSec; Sigma-Aldrich; See Table S3). Two different esiRNAs targeting CryAB and PFD2 were tetsted, of which only 1 resulted in knockdown and was analyze (HU-01337-3 for PFD2 and HU-01732-2 for CRYAB). The experiment was performed similarly, except that all six replicates for each esiRNA were performed in the same plate, and RSV-mKate2 was used. Virus replication was assessed at 24 hpi by quantifying the average red fluorescence in each well using a live cell microscope (Inncucyte S3, Sartorius).

### Evaluation of gene knockdown

A549 cells were reverse transfected as indicated above. Following 72 hours, gene expression was measured by qPCR. For this, RNA was extracted from A549 cells using RNAzol RT (Sigma-Aldrich) following the manufacturer protocol. The cDNA was then synthesized from 500 ng of RNA and used to evaluate gene expression by qPCR using the primers listed in Table S4 and the M-MuLV Reverse Transcriptase (NZYtech) in a QuantStudio3 machine using the PowerUp SYBR green qPCR master mix (Applied Biosystems). Knockdown efficiency was calculated by standardizing target gene expression to that of GAPDH in esiRNA transfected cells versus control-esiRNA transfected cells using the ΔΔCt method. For a few esiRNA, high expression levels were observed in knockdown cells and these were retested individually and the strongest gene knock-down level was taken as the knockdown value.

### Toxicity evaluation

To assess esiRNA toxicity, cells were transfected in triplicate as indicated above and incubated at 37°C for 72 hours. Subsequently, the medium was replaced by fresh DMEM containing a final concentration of 44μM resazurin. After a 1-hour incubation period, fluorescence was read using excitation/emission filters of 535/595nm using a microplate reader (Tecan Spark). Toxicity was compared by standardizing to the average signal obtained from cells transfected with the control esiRNA. To assess CB-5083 toxicity, cells were treated with increasing concentrations of the inhibitor for 24 hours before the addition of resazurin as indicated above.

### Drug treatments

Cells were treated with different concentrations of the VCP/p97 inhibitor CB-5083 for 2 hours. Subsequently, cells were infected at an MOI of 1 for 2 hours in the presence of the drug. Following infection, the virus inoculum was removed and fresh media containing the indicated inhibitor concentration was added. Twenty-four hours later, the supernatant was collected and used for measuring virus production via a plaque assay as indicated below.

### Analysis of VSV-G surface expression

Hela-H1 cells were transfected with a plasmid encoding the VSV-G gene (pMD2.G; Addgene #12259) using Lipofectamine 2000. Twenty-four hours later, 1μM CB-5083 or DMSO were added to the media for an additional 24 hours. Finally, cells were trypsinized, stained with a primary anti-VSV antibody (Cuevas, Durán-Moreno and Sanjuán, 2017) (kind gift of Dr. Rafael Sanjuán), followed by a FITC labeled secondary goat anti-mouse IgG (ThermoFisher Scientific) and analyzed on a FACSVerse (BD Biosciences).

### Statistical analysis

All statistical tests were performed with R (version 4.1.2) and graphs produced with the ggpubr (version 0.4.0) package. All tests were two-tailed, and the individual test employed is indicated in the main text. Q-values were obtained using the q-value package (version. 3.14). Statistical significant was set at p < 0.05 or q < 0.05.

## RESULTS

### RNAi screen to identify chaperones involved in the replication of RSV

In order to define which cytoplasmic chaperones are involved in the RSV replication cycle, we selected a list of cytoplasmic cellular molecular chaperones as well as key modulators of proteostasis that are expressed in the lung (see methods; Table S1). From this list, endoribonuclease-prepared siRNA (esiRNA) were commercially available for 114 (Table S2). These included a total of 37 chaperones genes from all major cytoplasmic chaperone families (Hsp70, Hsp90, TRiC/CCT, Prefoldin, and small heat shock proteins), 73 co-chaperones (e.g. Hsp40s, proteins containing TPR domain), a central player in the protein degradation machinery that cooperates with different chaperone systems (VCP/p97), and three genes from transcription factors involved in the cellular heat shock response (HSF; See Table S2).

To identify cytoplasmic chaperones relevant to RSV infection, we used the human lung adenocarcinoma cell line A549 as a model for the alveolar type II pulmonary epithelium. We first assessed the effects of chaperone depletion on cell viability. A549 cells were transfected with the different esiRNAs and cell viability was assessed three days following transfection using the Alamar Blue (resazurin reduction) assay (Fig. S1A; Table S2). Most esiRNA transfections resulted in low or no toxicity, with greater than two-thirds of genes maintaining cell viability above 70% relative to non-targeting control transfection (77 of 114 genes). Strong toxicity was only observed for 5% of esiRNAs, which reduced viability >50% (Table S2). We next evaluated the effect of ~60% of the esiRNAs (69 of the 114) for their knockdown efficiency using qPCR to measure target gene expression. A median reduction of 70% was observed, with greater than two-thirds (47/69) of the genes showing less than 50% expression, suggesting gene-specific knockdown is achieved using the esiRNA transfection (Fig. S1B; Table S2).

We next evaluated the effect of chaperone depletion on RSV replication (Fig. 1A). To this end, cells were transfected with esiRNA targeting the different chaperones or a control esiRNA targeting firefly luciferase. Following 48 hours, cells were infected with an RSV strain carrying a nano-luciferase reporter (RSV-nanoLuc) at a low multiplicity of infection for 24 hours. Finally, nanoLuc activity was measured to quantify virus entry and intracellular replication and the results were standardized to that of control transfected conditions. Hits were evaluated using the strictly standardized mean difference (SSMD) and its associated significance value following multiple testing correction (q value) (Zhang, 2011). SSMD provides a measure of effect size that takes into account variability in both experimental and control conditions, with positive values indicating increased replication and negative values indicating reduced replication. In total, 24 genes showing a knockdown efficiency of at least 20% had a statistically significant effect on RSV replication (q < 0.05) that was classified as at least moderate (absolute SSMD score of >1.28; Fig. 1B; Table S5). The effect on RSV replication was not significantly associated with viability (p = 0.13, Spearman’s ρ = −3.2), suggesting the results are likely to stem from specific effects on RSV replication rather than toxicity. Within this dataset, genes belonging to all major chaperone systems were observed (e.g.Hsp70/HSPA, Hsp90/HSPC, TRiC/CCT, prefoldin), the multifunctional VCP/p97 protein, master regulators of the transcriptional response to stress (Heat Shock Factor; HSF1), and co-chaperones regulating the activity of Hsp70 and Hsp90 (Fig. 1B; Table S5). Nearly all of the identified chaperones resulted in reduced replication following their depletion, indicating a supportive role in the RSV replication cycle; however, the replication of RSV increased following the depletion of two chaperones, SGTB and DNAJC12, suggesting these could be detrimental to RSV replication under normal conditions (Fig. 1B; Table S5). Importantly, we identify isoforms of both the Hsp70 and Hsp90 chaperones to inhibit RSV replication, as has been previously reported using pharmacological inhibitors (Geller, Andino and Frydman, 2013; Munday *et al*., 2015a; Latorre, Mattenberger and Geller, 2018), confirming the ability of the screen to identify relevant chaperones.

**Figure 1.**
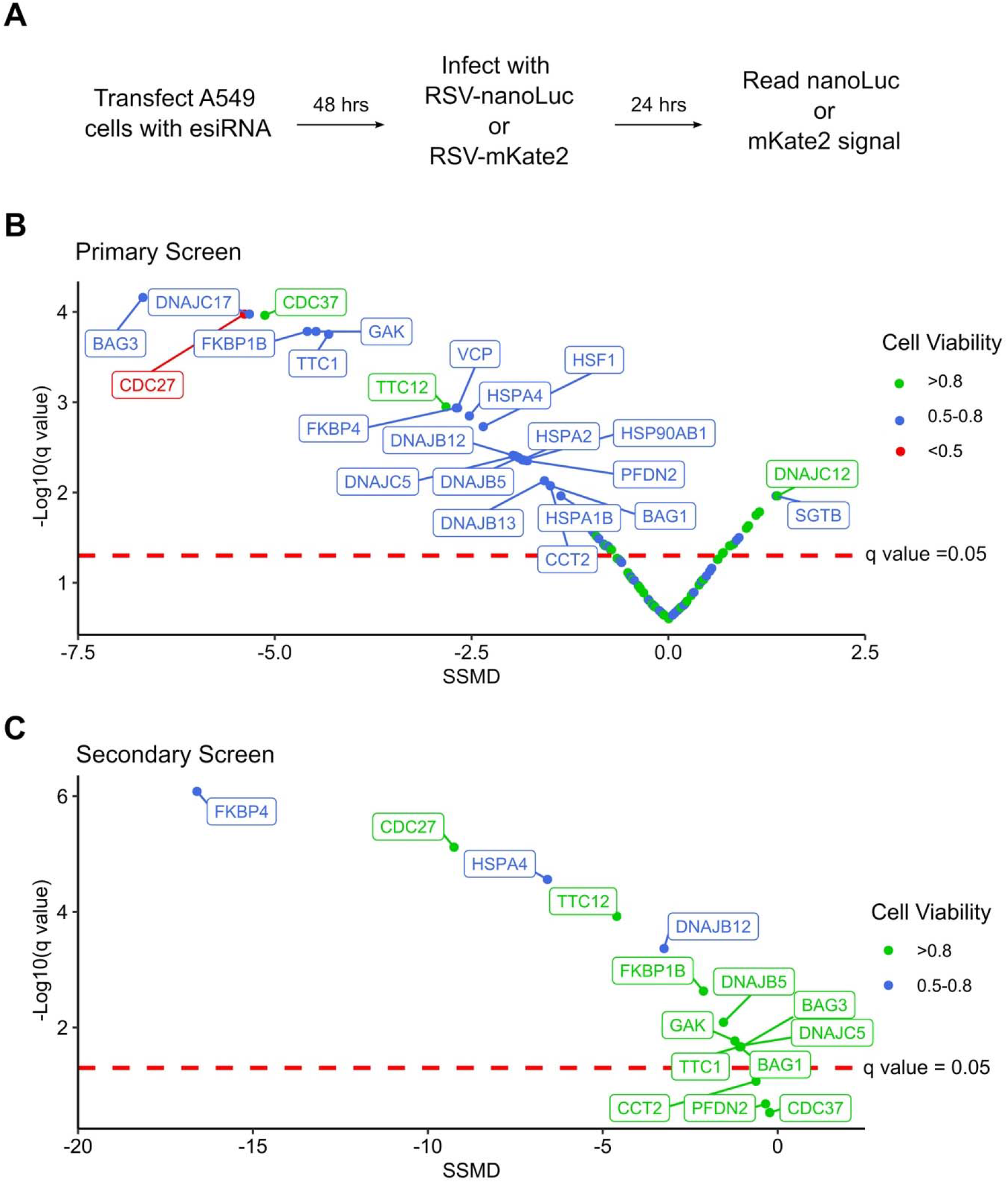
An RNAi screen for chaperones and co-chaperones involved in RSV replication. **A**. Overview of the experimental approach. **B**. Results of the primary screening, showing the strictly standardized mean deviation (SSMD) and the statistical significance (q-value). n=6. **C**. Results of the secondary screening, showing the SSMD and statistical significance. n=6.

To validate the results of the primary screen, we obtained esiRNA targeting alternative sites in 15 genes that showed a significant reduction in RSV replication when depleted (Table S5). Moreover, to account for any effects the chaperones could have on the nanoLuc reporter rather than RSV replication, we used an RSV strain encoding a red fluorescent gene (RSV-mKate2) for the validation test. From the 15 genes analysed, 80% significantly reduced RSV replication (q < 0.05; 12/15 genes) without strong toxicity (viability >70%; Fig. 1C and Table S5) supporting the validity of the primary screen. For two of the three genes that did not show a significant effect on replication in the secondary screen, lower knockdown levels were observed (~2-fold and 6-fold lower for CCT2 and PFDN2, respectively), which could underlie the lack of effect. For the third gene, CDC37, a similar reduction in gene expression was observed in both the primary and secondary screens, indicating this gene is not likely to represent a real hit. A very strong reduction in RSV replication was observed for FKBP4, a cis-trans prolyl isomerase from the immunophilin family that interacts with Hsp70 and Hsp90, the Hsp70 nucleotide exchange factor HSPA4, and the Hsc70 co-chaperone DNAJB12 (Fig. 1C; Table S5).

### VCP/p97 inhibition reduces the replication of RSV and vesicular stomatitis virus

Pharmacological inhibitors of cellular factors provide a direct means to assess the relevance of the targeted protein in viral infection. Indeed, Hsp70 and Hsp90 inhibitors have been used to demonstrate the relevance of these chaperones in the viral replication cycle (Geller, Taguwa and Frydman, 2012; Geller, Andino and Frydman, 2013; Aviner and Frydman, 2019; Yang *et al*., 2020). We therefore chose to validate the antiviral role of VCP/p97, which was identified in the primary screen (Figure 1B), using a selective inhibitor that prevents ATP binding to VCP/p97, CB-5083 (Zhou *et al*., 2015; Huryn, Kornfilt and Wipf, 2019). We first evaluated the toxicity of CB-5083 in A549 cells. Concentrations of up to 0.5μM resulted in low toxicity (~90% viability), while higher concentrations resulted in significant toxicity (~50% viability at 2μM; Fig. 2A). Next, we assessed the effect of VCP/p97 inhibition on RSV production. For this, cells were infected in the presence of VCP/p97 and RSV production quantified following 24 hours. At CB-5083 concentrations of 0.5μM, RSV production was reduced by ~90% (p < 0.01 by Mann-Whitney test), while concentrations of 1μM resulted in a >99% reduction but showed higher toxicity (p<0.01 by Mann-Whitney test; Fig. 2A,B). To ensure the effect was not cell-specific, we repeated the experiments in Hela-H1 cells. At concentrations that did not alter cell viability strongly (>92% viability at 0.5μM; Fig. 2C), RSV production was reduced by 84% (p<0.05 by Mann-Whitney test; Fig. 2D). Interestingly, VCP/p97 inhibition also reduced the production of an unrelated member of the *Mononegavirales* order, vesicular stomatitis virus (VSV), by 98% at the same concentration (p<0.05 by Mann-Whitney test; Fig. 2E). Inhibitor concentrations showing higher toxicity (25% toxicity; 1μM) had profound antiviral activity, reducing RSV and VSV production by >3 and >5 logs, respectively (p<0.01 and p<0.001 by Mann-Whiney test for RSV and VSV, respectively; Fig. 2D & Fig. 2E). Importantly, these antiviral effects are not the result of toxicity, as the replication of a positive-strand RNA virus, coxsackievirus B3 (CVB3), was not affected at inhibitor concentrations up to 1μM (p>0.05 by Mann-Whitney test; Fig. 2F), as has been previously reported (Arita, Wakita and Shimizu, 2012).

**Figure 2.**
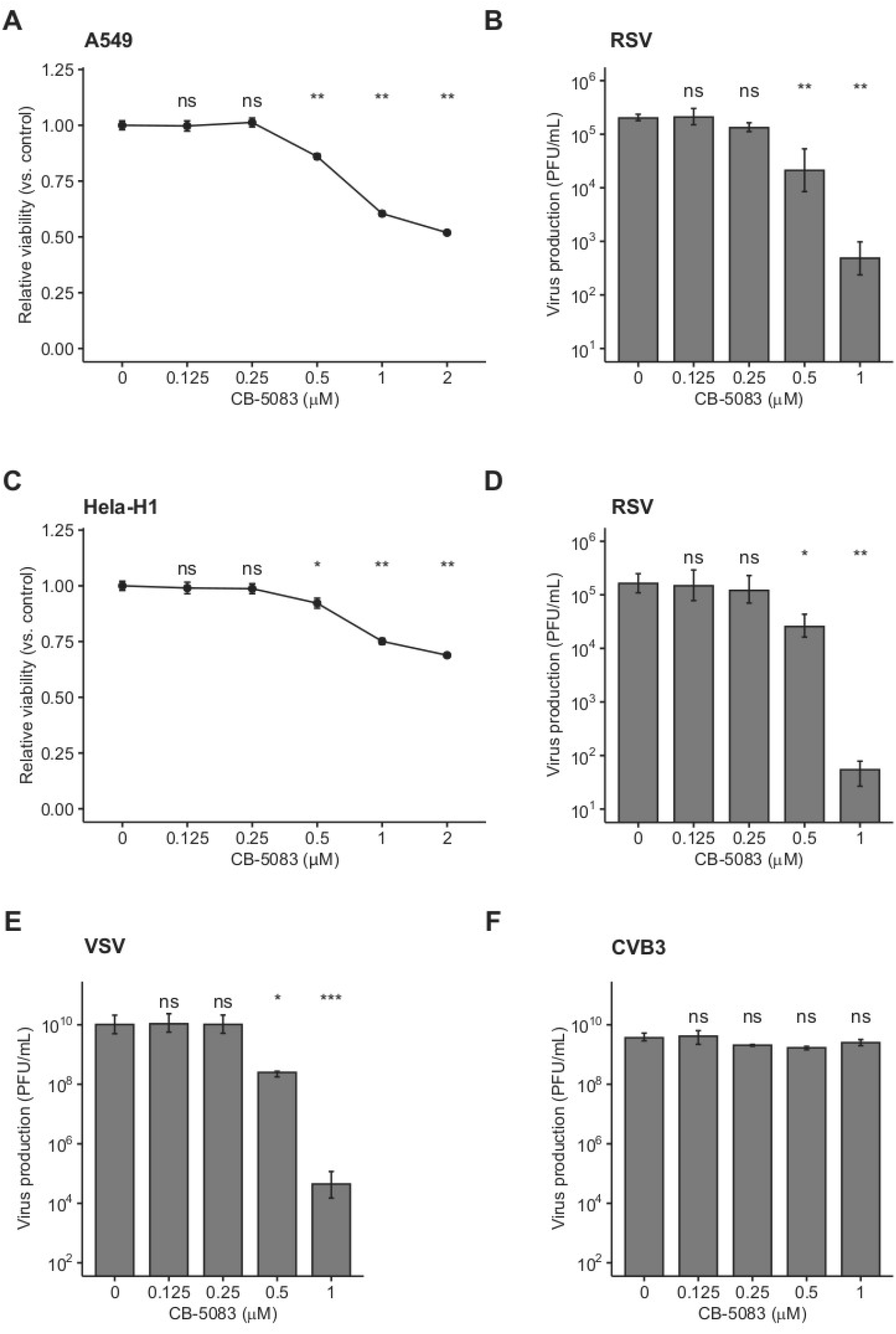
VCP/p97 inhibition reduces RSV and VSV production. **A**. The effect of the VCP/p97 inhibitor CB-5083 on A549 cell viability. **B**. The effect of CB-5083 on RSV virus production in A549. **C**. The effect of the VCP/p97 inhibitor CB-5083 on Hela-H1 cell viability. **D-E**. The effect of CB-5083 on RSV (**D**), vesicular stomatitis virus (VSV) (**E**), or coxsackievirus B3 (CVB3) production. (**F**) virus production in Hela-H1 cells. Results indicate the mean and SEM of >3 replicates. ns, p>0.05; **, p<0.01; ***, p<0.001; ****, p<0.0001 by Mann-Whitney test.

### VCP/p97 is not involved in the early steps of viral infection

To determine whether VCP/p97 is involved in the early steps of viral replication, namely entry and uncoating, we performed a time-of-addition study. Specifically, we added the VCP/p97 inhibitor CB-5083 at 1μM to HeLa-H1 cells either 1 hour before infection, following infection, or 2 hours postinfection, when binding, entry, and uncoating are complete. We then used the fluorescence intensity of the viral-encoded reporter proteins to measure effects on viral replication. For RSV we observed an 87%, 82%, and 79% reduction in fluorescence intensity when the drug was added 1 hour before infection, at the time of infection, or 2 hours post-infection, respectively, compared with the DMSO-treated cells (p < 0.01 for all versus DMSO by t-test on log-transformed data; Fig. 3A). Similarly, VSV replication was reduced by 99.7%, 99.8%, and 99.9% when added 1 hour before infection, at the time of infection, or 2 hours post-infection, respectively, compared with the DMSO-treated cells (p < 0.01 for all conditions versus DMSO by t-test on log-transformed data; Fig. 3B), with no appreciable reduction in effectivity when the drug was added at a stage following entry and uncoating. As the effect of VCP/p97 inhibition on the replication of both RSV and VSV was similar when added prior to infection or after the completion of entry, VCP/p97 likely acts downstream of these early steps in the viral infection cycle.

**Figure 3.**
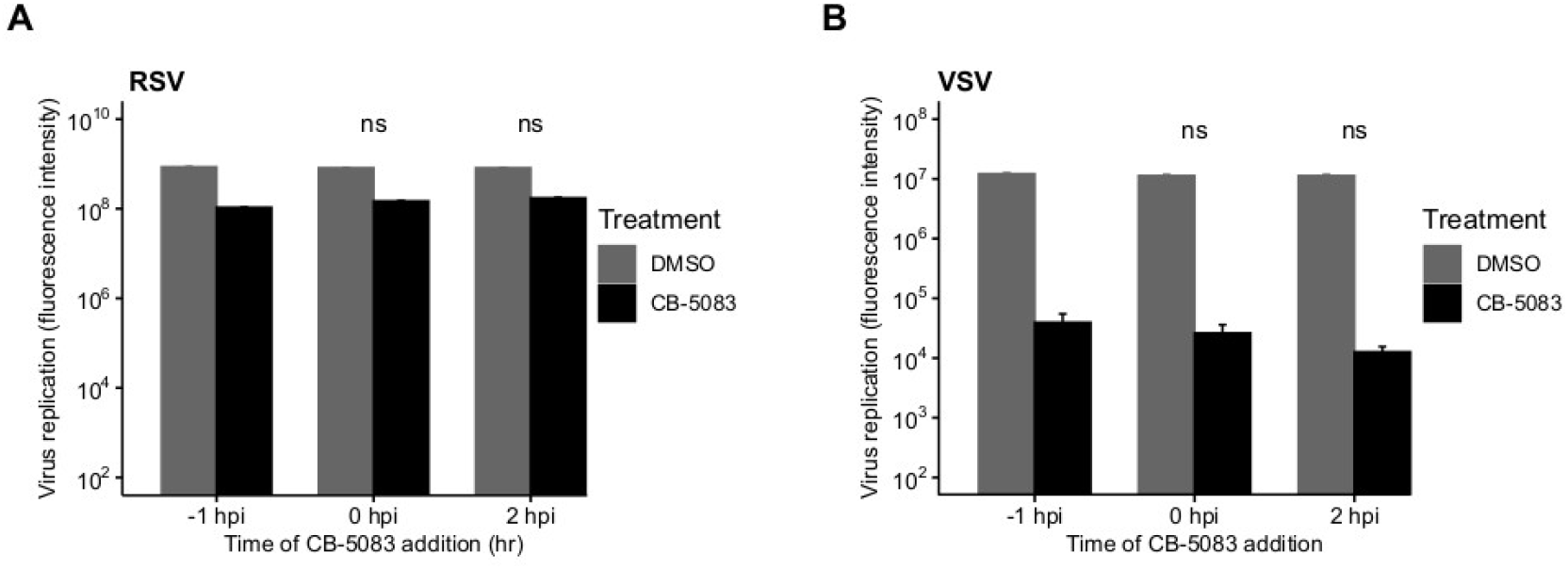
VCP/p97 is not involved in the early steps of infection. The VCP/p97 inhibitor CB-5083 was added at the indicated time point relative to infection (hours post-infection, hpi) with RSV (**A**) or VSV (**B**) and virus production was quantified. Data represent mean and SEM with an n=3. ns, p>0.05 by t-test on log-transformed data.

### VCP/p97 is involved in multiple steps of the viral replication cycle

The final steps in the viral replication cycle are the formation of mature virions and egress from the cell. To investigate whether VCP/p97 plays a role in these late stages of viral replication, we evaluated how CB-5083 affected intracellular viral replication compared to infectious particle formation in HeLa-H1. We reasoned that observing a stronger effect on the quantity of released infectious particles compared to intracellular replication will support an additional role for VCP/p97 in the late stages of viral replication. As a control, we tested the effects of an Hsp90 inhibitor, 17-DMAG (Egorin *et al*., 2002), under the same conditions. Hsp90 is known to be required for the folding and maturation of the RNA-dependent RNA polymerase of both RSV (Geller, Andino and Frydman, 2013) and VSV (Connor *et al*., 2007). As polymerase function is required for replication but is not involved in late stages of the viral cycle, blocking Hsp90 should show a similar effect on both intracellular replication and virus production.

As in previous experiments (Fig. 2B,D), both RSV intracellular viral replication, as measured by the quantification of viral encoded fluorescent protein, and virus production were significantly reduced by VCP/p97 inhibition at 1μM (Fig. 4A, B). However, virus production was reduced by ~4-logs (p < 0.0001 by t-test on log-transformed data; Fig. 4A), while intracellular replication was only reduced by ~1-log when blocking VCP/p97 function (p < 0.001 by t-test on log-transformed data; Fig. 4B). When standardizing the effect of VCP/p97 inhibition in each assay to control-treated cells, virus production was >1,000 times more affected by VCP/p97 inhibition than intracellular replication (p < 0.01 by t-test on log-transformed data; Fig. 4C). In contrast, while Hsp90 inhibition also reduced both RSV virus production (p < 0.001 by t-test on log-transformed data; Fig. 4A) and intracellular viral replication (p < 0.001 by t-test on log-transformed data; Fig. 4B), the relative effect on each one was similar (average standardized effect of intracellular replication versus virus production of 3.44 ± 1.12; p <0.05 by t-test on log-transformed data; Fig. 4C). This is consistent with no additional role of the Hsp90-dependent viral RNA-dependent RNA polymerase in the late stages of replication. Similarly, for VSV, CB-5083 treatment diminished virus production by ~6 logs (p < 0.0001 by t-test on log-transformed data; Fig. 4D) and intracellular replication by ~3 logs respectively (p < 0.0001 by t-test on log-transformed data; Fig. 4E). As for RSV, virus production was >1,000 times more affected by VCP/p97 inhibition than intracellular replication (p<0.0001 by t-test on log-transformed data; Fig. 4F). Hsp90 inhibition showed only mild inhibition of VSV virus production in our assay conditions, with a 3-fold reduction observed for virus production (p < 0.01 by t-test on log-transformed data; Fig. 4C), and the effect on intracellular replication was not significant (p > 0.05 by t-test on log-transformed data; Fig. 4D), potentially due to the late time point of infection at which we examined intracellular replication (24 hours). Again, Hsp90 inhibition affected both VSV virus production and intracellular replication in a similar manner, indicating no specific role in late events (average standardized effect of intracellular replication versus virus production of 3.47 ± 0.73; p <0.01 by t-test on log-transformed data; Fig. 4F)

**Figure 4.**
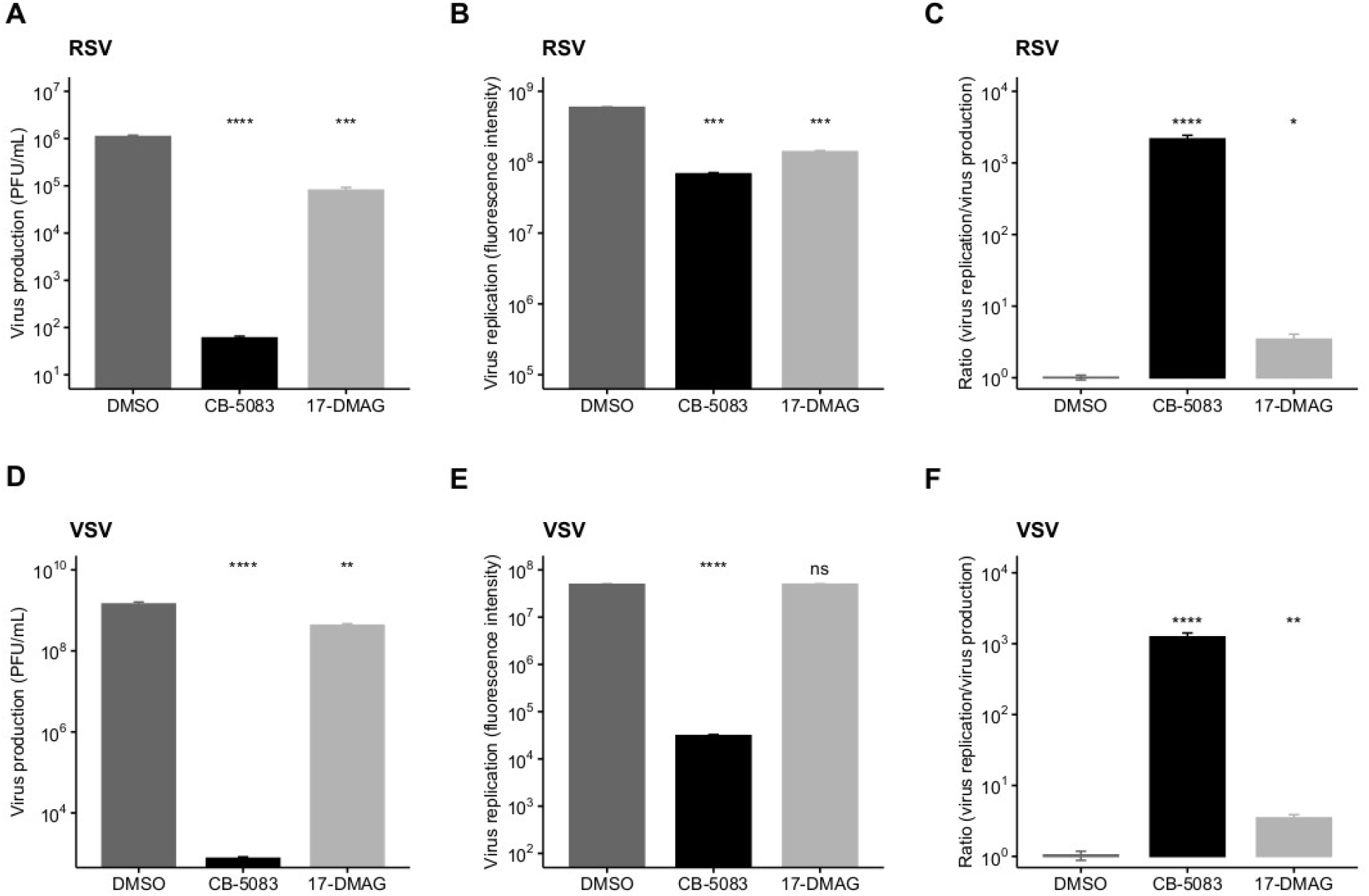
VCP/p97 is involved in multiple steps of the RSV and VSV infection cycle. Cells were treated with the VCP/p97 inhibitor CB-5083 or the Hsp90 inhibitor 17-DMAG and infected with RSV (top row) or VSV (bottom row). Subsequently, virus production was measured by plaque assay (**A,D**) and intracellular replication was measured by the expression of a reporter protein (**B,E**). The ratio of the antiviral effect observed for virus replication compared to virus production (**C,F**). Plotted are the means and SEM for three replicates. ns, p>0.05; **, p<0.01; ***, p<0.001; ****, p<0.0001 by t-test on log-transformed data.

### VCP/p97 inhibition does not affect trafficking to the cell surface

Both RSV and VSV encode glycoproteins that require translation, folding, and trafficking in the endoplasmic reticulum (ER). Since VCP/p97 is a key player in ER quality control and ER-associated degradation (van den Boom and Meyer, 2018; Hänzelmann, Galgenmüller and Schindelin, 2019; Huryn, Kornfilt and Wipf, 2019), we examined whether the translation of the VSV glycoprotein (VSV-G) was affected by VCP/p97 inhibition. For this, cells were transfected with a plasmid encoding VSV-G and subsequently treated with CB-5083 (1μM) or DMSO for 24 hours prior to assessing VSV-G surface expression by flow cytometry. Surprisingly, CB-5083 treatment resulted in a slightly enhanced expression of the VSV-G protein on the cell surface (p < 0.01 by t-test; Fig. 5). Hence, the antiviral activity of VCP/p97 inhibition is not due to reduced translation or trafficking in the ER. However, it is important to note that these experiments do not assess whether the VSV-G protein is properly folded and functional.

**Figure 5.**
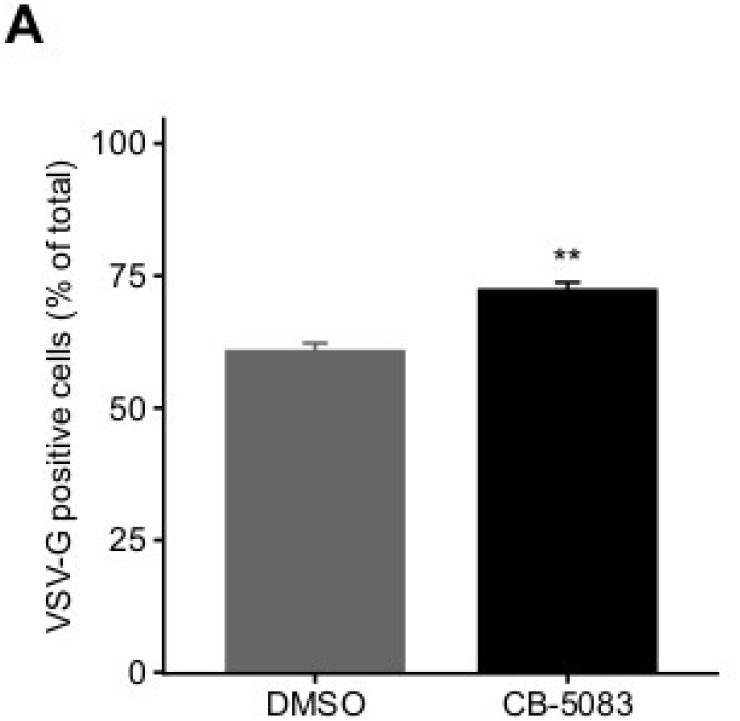
VCP/p97 treatment does not block the trafficking of viral glycoproteins to the surface. Cells were transfected with a plasmid encoding the VSV glycoprotein (VSV-G). Following 24 hours, cells were either treated or mock-treated with the VCP/p97 inhibitor CB-5083. Following an additional 24 hours, VSV-G expression was assessed by flow cytometry. Data represent the mean and SEM of 3 replicates. **, p<0.01 by t-test.

## DISCUSSION

Cellular proteostasis is maintained through highly interconnected networks of chaperones, cochaperones, and components of the degradation machinery (Kim *et al*., 2013; Balchin, Hayer-Hartl and Hartl, 2016; Bar-Lavan, Shemesh and Ben-Zvi, 2016). The complexity of these interactions makes it challenging to define cellular proteostasis networks for particular proteins and/or cellular functions. As RNA viruses utilize the cellular proteostasis machinery but encode only few proteins, they could provide a simplified system with which to define such proteostasis networks.

To date, very limited information is available on the cellular proteostasis machinery involved in RSV replication. Prior work has demonstrated a dependence on both Hsp70 and Hsp90 chaperone systems (Radhakrishnan *et al*., 2010; Geller, Andino and Frydman, 2013; Munday *et al*., 2015a; Latorre, Mattenberger and Geller, 2018), largely for polymerase function, but has not defined which isoform of these chaperones is of relevance, nor the co-chaperones that are involved. In this work, we set out to begin to define the cellular protein folding networks utilized by RSV during replication. We assessed a total of 114 genes belonging to all major chaperone systems using an RNAi screen and identified a total of 24 cellular genes that significantly impacted RSV replication, having at least a moderate effect (absolute SSMD > 1.28; Fig. 1B). To validate the primary screening, we used RNAi targeting a different site in 15 hits from the primary screen. Overall, the knockdown of 12 genes reduced RSV replication significantly (Fig. 1C). For two of the three genes that did not reach significance (CCT2 and PFD2), reduced levels of gene knockdown efficiency relative to the primary screen could underlie the lack of effect (Table S5). Finally, we validated the role of one of the genes identified in the primary screen, VCP/p97, using a pharmacological approach (Fig. 2). Hence, of the 24 genes identified in the primary screening, we validated 12/15 by RNAi, one by pharmacological inhibition (VCP/p97), and three were implicated in previous studies (two Hsp70 isoforms and one Hsp90 isoform), suggesting >80% of the genes are likely to represent true hits.

Overall, numerous genes involved in the Hsp70 chaperone cycle were identified to alter RSV replication with at least a moderate effect (absolute SSMD > 1.28): two of nine Hsp70 chaperones (HSPA1B and HSPA2), three of nine Hsp70 nucleotide exchange factors (BAG1, BAG3, and HSPA4), and seven of the 24 Hsp70 co-chaperones (Table S5). Previously, the constitutive Hsp70 isoform, Hsc70/HSPA8, was shown to be involved in RSV replication (Radhakrishnan *et al*., 2010). Indeed, we observe this gene to significantly reduce RSV replication in our screen (q-value<0.05) although its effect was relatively weak (SSMD = −0.79), as was the case for two additional Hsp70 isoforms (HSPA1A and HSPA12B; Table S2). Overall, our results reduce the complexity of the Hsp70 system to 12 genes of the 42 genes analysed. Similarly, we identified one of the two Hsp90 isoforms (HSP90AB1) and two of the fourteen Hsp90 co-chaperones analysed (FKBP1B and FKBP4) to be involved in the replication of RSV, reducing the members of this chaperone family to three out of 16 genes. Moreover, four of nineteen Hsp70-Hsp90 co-chaperones containing TPR domains were identified (CDC27, TTC1, TTC12, and SGTB). These results suggest that despite numerous chaperones and co-chaperones in each family, individual proteins have specific functions that cannot be compensated by other isoforms. This is in agreement with the results of previous work characterizing the role of different components of the Hsp70 chaperone system in the replication of Dengue and Zika virus (Taguwa *et al*., 2015, 2019). The results of the current work can now facilitate definition of specific chaperone subnetworks involved in individual steps of the RSV infection cycle.

Previous work has shown that pharmacological inhibition of Hsp70 and Hsp90 can constitute a broad-spectrum antiviral approach (Geller *et al*., 2007; Geller, Taguwa and Frydman, 2012; Latorre, Mattenberger and Geller, 2018; Aviner and Frydman, 2019). As a clinically evaluated pharmacological inhibitor (CB-5083) was available for one of the identified hits from the RNAi screen, VCP, we evaluated its potential as an antiviral. We found CB-5083 to inhibit RSV replication in both A549 and Hela-H1 cells (Fig. 2B,D), as well as to block the replication of VSV (Fig. 2E). These results further support VCP/p97 as a general target for antiviral therapy against different viruses (Das and Dudley, 2021). To gain insights into the step of the replication cycle where VCP/p97 acts, we examined the effect of adding CB-5083 before or after viral entry. We found no difference in the antiviral effect of CB-5083 if added post-entry, suggesting a role in downstream aspects of viral replication (Fig. 3). In contrast, the effect of CB-5083 was observed for both intracellular replication and virus release, with the latter being significantly more affected (Fig. 4). Nevertheless, this is unlikely to stem from effects on viral glycoprotein expression, as blocking of VCP/p97 did not reduce the expression of the VSV glycoprotein (Fig. 5). Hence, it is likely that VCP/p97 acts at multiple steps of the replication of RSV and VSV.

## Author Contributions

VL performed the work and analyzed the data. RG performed the work, analyzed the data, obtained funding, and wrote the manuscript.

## Acknowledgments

We would like to thank Dr. Beatriz Alvarez-Rodriguez for critical reading of the manuscript.

## SUPPLEMENTARY DATA

**Figure S1.**
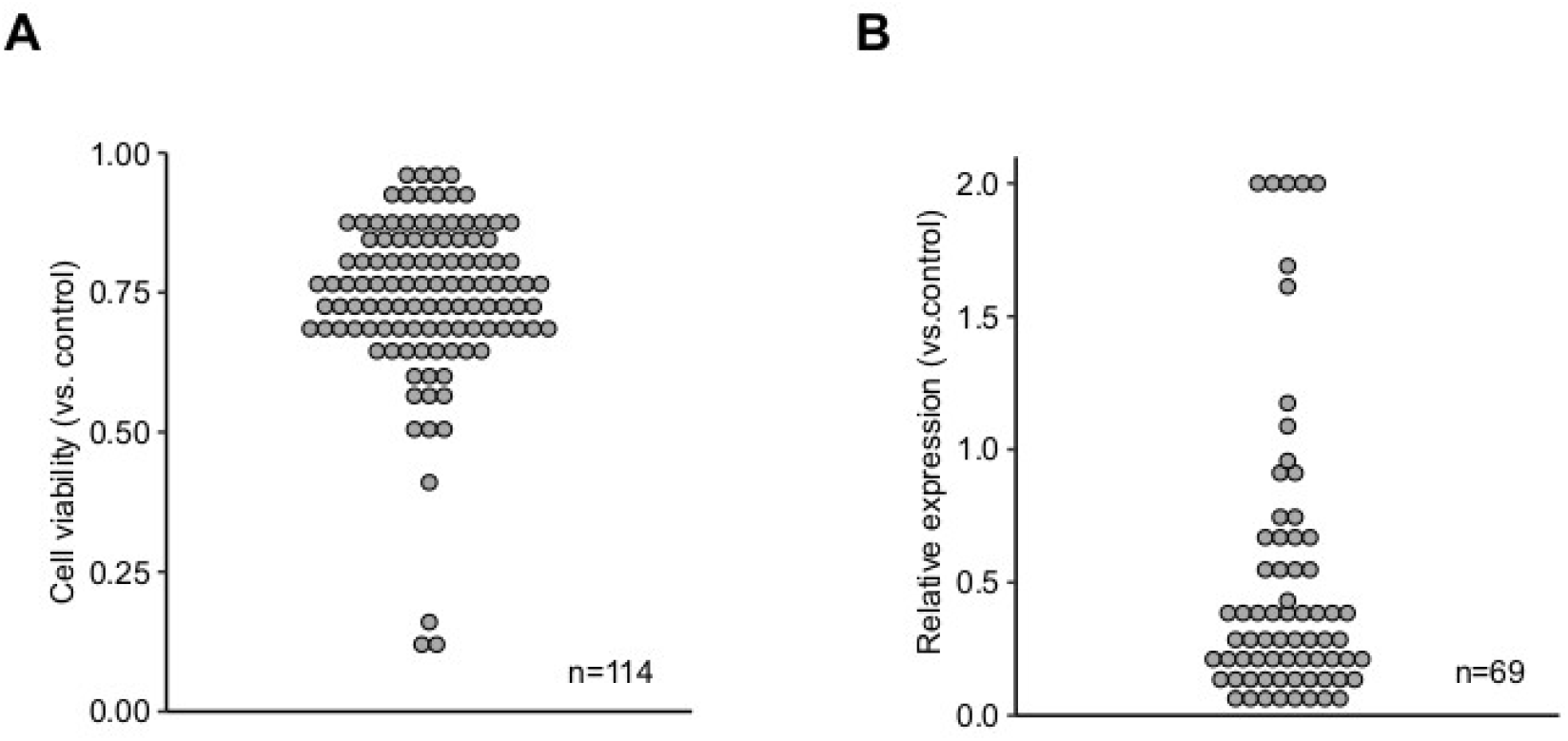
Evaluation of the toxicity and knockdown efficiency for the esiRNAs used in the primary screen. **A**. The relative viability of cells transfected with esiRNAs targeting cellular proteostasis components was compared to that of cells transfected with non-targeting esiRNA. **B**. The expression of the targeted genes following esiRNA transfection was compared to that in cells transfected with nontargeting esiRNA.

## SUPPLEMENTARY TABLES

**Table S1:** Description of the genes selected for evaluation

**Table S2:** Results of the toxicity, knockdown efficiency, and SSMD for evaluated genes

**Table S3:** esiRNA information for screened genes

**Table S4:** Primer sequences for evaluation of gene knockdown using qPCR

**Table S5:** Significant hits from the primary and secondary screen RNAi screens

